# Natural selection in the evolution of SARS-CoV-2 in bats, not humans, created a highly capable human pathogen

**DOI:** 10.1101/2020.05.28.122366

**Authors:** Oscar A. MacLean, Spyros Lytras, Steven Weaver, Joshua B. Singer, Maciej F. Boni, Philippe Lemey, Sergei L. Kosakovsky Pond, David L. Robertson

## Abstract

RNA viruses are proficient at switching host species, and evolving adaptations to exploit the new host’s cells efficiently. Surprisingly, SARS-CoV-2 has apparently required no significant adaptation to humans since the start of the COVID-19 pandemic, with no observed selective sweeps since genome sampling began. Here we assess the types of natural selection taking place in *Sarbecoviruses* in horseshoe bats versus SARS-CoV-2 evolution in humans. While there is moderate evidence of diversifying positive selection in SARS-CoV-2 in humans, it is limited to the early phase of the pandemic, and purifying selection is much weaker in SARS-CoV-2 than in related bat *Sarbecoviruses*. In contrast, our analysis detects significant positive episodic diversifying selection acting on the bat virus lineage SARS-CoV-2 emerged from, accompanied by an adaptive depletion in CpG composition presumed to be linked to the action of antiviral mechanisms in ancestral hosts. The closest bat virus to SARS-CoV-2, RmYN02 (sharing an ancestor ∼1976), is a recombinant with a structure that includes differential CpG content in Spike; clear evidence of coinfection and evolution in bats without involvement of other species. Collectively our results demonstrate the progenitor of SARS-CoV-2 was capable of near immediate human-human transmission as a consequence of its adaptive evolutionary history in bats, not humans.

## Introduction

In December 2019 an outbreak of pneumonia cases in the city of Wuhan, China, was linked to a novel coronavirus. Evolutionary analysis placed this new virus to humans in the same subgenus of *Betacoronaviruses*, the *Sarbecoviruses*^1^, that the SARS virus belongs to, and it was named SARS-CoV-2 to reflect its status as a sister lineage to the first SARS virus^2^. This novel lineage represents the seventh known human-infecting member of the *Coronaviridae*. The initial outbreak of human cases of the virus was connected to the Huanan Seafood Wholesale Market in Wuhan^3^, and while related viruses have been found in horseshoe bats^4^ and pangolins^5^, their divergence represents decades of evolution^6^ leaving the direct origin of the pandemic unknown. In addition to elucidating the transmission route from animals to humans, key questions for assessing future risk of emergence are: (i) what is the extent of evolution required for a bat virus to transmit to humans, and (ii) what subsequent evolution must occur for efficient transmission once the virus is established within the human population?

The first SARS virus outbreak in 2002/2003, causing approximately 8,000 infections, and its re-emergence in late 2003, causing four infections, were linked to Himalayan palm civets and raccoon dogs in marketplaces in Guangdong province^7,8^. Later it became clear that these animals had been conduits for spillover to humans and not true viral reservoirs^9^. Extensive surveillance work subsequently identified related viruses circulating in horseshoe bats in China, some of which can replicate in human cells^10,11^. The bat viruses most closely related to SARS-CoV-1, can bind to human angiotensin converting enzyme (ACE2, the receptor SARS-CoV-2 also uses for cell entry), while the addition of protease is required for the more divergent bat viruses tested (red and grey lineages respectively, Supplementary Figure 1)^12^. Collectively these results demonstrate that, unlike other RNA viruses assumed to acquire adaptations after switching to a new host species for efficient replication and/or spreading, the *Sarbecoviruses* – which already transmit frequently among bat species^13^ – can exploit the generalist properties of their ACE2 binding ability, facilitating successful infection of non-bat species, including humans. A main difference between SARS-CoV-1 and −2 is the increased binding affinity for human ACE2^14^ permitting more efficient use of human cells and the upper respiratory tract, and on average lower severity but, paradoxically – due to greater penetrance into the human population – higher disease burden.

There is intense interest in the mutations emerging in the SARS-CoV-2 pandemic^15–17^. Although the vast majority of observed genomic change is expected to be ‘neutral’^18,19^, mutations with functional significance to the virus will likely arise, as they have in many other viral epidemics and pandemics^20^. In SARS-CoV-2 amino acid replacements in the Spike protein could reduce the efficacy of vaccines, replacements in proteases and polymerases could result in acquired drug resistance, and other mutations could change the biology of the virus, e.g., enhancing its transmissibility or severity^21^, contributing to adaption to us, a new host species. A main way to begin to understand the functional impact of mutations is to characterise the selective regime they are under. Mutations which are under positive selection are of particular interest as they are more likely to reflect a functional change. However, identifying mutations under positive selection from frequency data alone can be misleading, as allele frequencies in viral pandemics are significantly driven by biased sampling, founder effects and superspreading events^17^. Exponentially growing populations can increase in average fitness^22^, however they are also expected to exhibit elevated genetic drift, with deleterious mutations surfing expansion waves^23^. Here, we investigate the ability of bat *Sarbecoviruses* to readily infect non-bat species through a comprehensive search for signatures of positive selection (a measure of molecular adaptation) in the virus circulating in humans since the COVID-19 outbreak began, and contrast this to historic selection acting on related bat viruses.

### Evidence of nearly neutral evolution, and relatively weak purifying selection in SARS-CoV-2

We first analyse selection acting on the encoded amino acids of SARS-CoV-2 using 50,240 QC filtered genome sequences from the GISAID database as of the 28th of June 2020, representing a sample of the variants circulating in humans (see Methods). Purifying selection acts more strongly on nonsynonymous sites, supported by the estimate that the nonsynonymous substitution rate (dN) was only 4% of speed of the synonymous substitution rate (dS) in the divergence between SARS-CoV-2 and bat virus RaTG13^24^. We used the phylogenetically corrected SLAC counting method^25^ to calculate “frequency-stratified” dN/dS estimates. This qualitative analysis revealed that, on average, the evolutionary process is weakly purifying, with point estimates of dN/dS ranging from 0.6 to 1.0, and generally declining with increasing frequency, with the exception of the high frequency variants (seen in more than 3,000 sequences), where dN/dS is 1.4, indicating weak positive selection (Figure 1A). The vast majority of observed mutations occur at low frequency, with 90% of mutations observed in fewer than 15 of the 50,240 sequences (Figure 1A), consistent with a model of exponential population growth of virus spread (Supplementary Figure 2, Supplementary Text 1). Initial searches for signatures of non-neutral selection in the SARS-CoV-2 sequence data from late March produced numerous false positive signals, due to artefactual lab recombination and other sequencing errors found on terminal branches (Supplementary Text 2-3).

**Figure 1.**
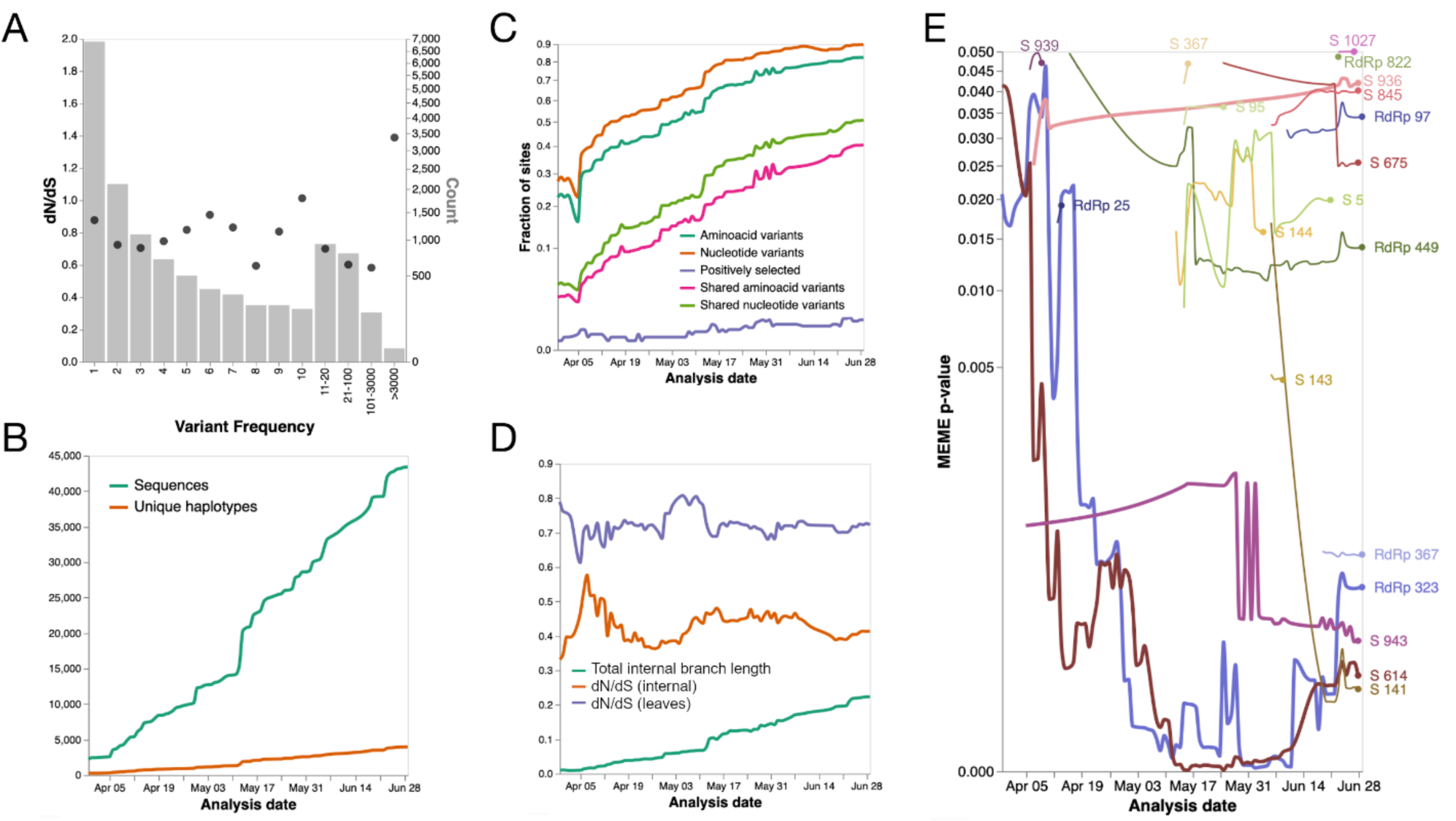
**(A)** Estimates of molecular adaptation (dN/dS) for 50,240 SARS-CoV-2 genome sequences based on the counting SLAC method^25^ (black circles) and the number of variants as a function of their frequency (grey bars). **(B)** Cumulative number of Spike gene sequences in GISAID passing QC filters from 5^th^ April to 28^th^ June, and the number unique haplotypes among them. **(C)** Cumulative fractions of codon sites in Spike which harbour different types of sequence variants or are positively selected (MEME^31^, internal branches only). **(D)** Estimates of gene-wide dN/dS in Spike on internal branches and terminal branches (MG94xREV model), and the total length of internal tree branches, which serves as a good proxy of statistical power to detect selection. And, **(E)** Statistical evidence for episodic positive selection in codons in SARS-CoV-2 S and RdRp that reached significance (p≤0.05) at least once during the analysis period. Four sites that appeared before April 15th and persisted are shown with thicker lines.

Subsequently, we have been performing daily analyses of the data excluding these terminal branches (http://covid19.datamonkey.org). As more sequences are deposited in public databases (Figure 1B), we expect to see increasingly more variation, with 89.6% of codons in Spike (S) containing some variation in most recent analyses (Figure 1C). Similar increasing trends are seen for sites with amino-acid variants (82.2%) and variants shared by more than one sequence (40.4% for amino-acids). However, the number of unique sequences has increased more slowly than the number of sequenced genomes (Figure 1B), and the fraction of sites inferred to be under positive selection remains below 1%. This observation shows SARS-CoV-2 is evolving relatively slowly with no dramatic increases in selective pressures occurring over this period. This pattern is further confirmed by the relatively stable behavior of gene-wide dN/dS estimates (Figure 1D), both on internal branches, which include successful transmissions, and terminal branches, which sometimes include intra-host “dead-ends” or deleterious variation^26^.

Even in Spike, which is being assiduously scrutinized for selection due to its immunogenic and phenotypic importance, overall selective pressure is stable over time, and consistent with weak purifying selection. The genetic homogeneity of SARS-CoV-2 results in very shallow phylogenetic trees, despite 10,000s of collected sequences, with cumulative branch lengths only about 0.2 substitutions/site (Figure 1D), somewhat limiting the statistical power of comparative methods^27^. There are individual sites in the SARS-CoV-2 genomes which possess statistically detectable signatures of episodic diversifying selection along internal tree branches. In S and RdRp, there are four sites that appear to be selected in data prior to April 15 (early pandemic, Figure 1E). These include Spike 614 and RdRp 323 which are in nearly perfect linkage disequilibrium (R^2^>0.99), with the former extensively discussed by others as having some functional significance^28^, while the latter received much less attention. The majority of these sites, however, are not detected until later in the pandemic, and most are transientis because mildly deleterious mutations in viral outbreaks fail to persist over longer time periods, being gradually purged by purifying selection^26,30^. Exponential growth, consistent with the SARS-CoV-2 trajectory (Supplementary Figure 2), is known to reduce the efficacy of purifying selection^23^, with deleterious mutations able to surf expansion waves. It is likely that many of these putatively deleterious segregating mutations will ultimately be lost, reducing the long term dN/dS, though their persistence will be influenced by future demographic patterns^23^, and the effect of lockdowns on the circulating variants.

### What about the bats? Positive selection in *Sarbecoviruses*

Coronaviruses frequently recombine in their bat hosts, with the Spike open reading frame (ORF) being an apparent hotspot for this process, with potentially adaptive implications for the viruses, i.e., antigenic shift, in the context of immune evasion^9,32–35^. To avoid the confounding effects of recombination on the inference of selection patterns, we separately analysed putatively non-recombinant regions, as derived in Boni et al. (2020; see Methods)^6^. For each region, we define as the ‘nCoV clade’ the set of viruses closest to SARS-CoV-2 in the phylogeny (see Methods). We find that genomic sites are generally subject to conservation in this nCoV clade, with 8184/9744 (84%) of codon sites conserved at the amino-acid level, and 4274 (43.7%) sites, of which 3388 were variable at the nucleotide level, showing evidence of purifying selection in this lineage (using the FEL method^25^; Figure 2).

**Figure 2.**
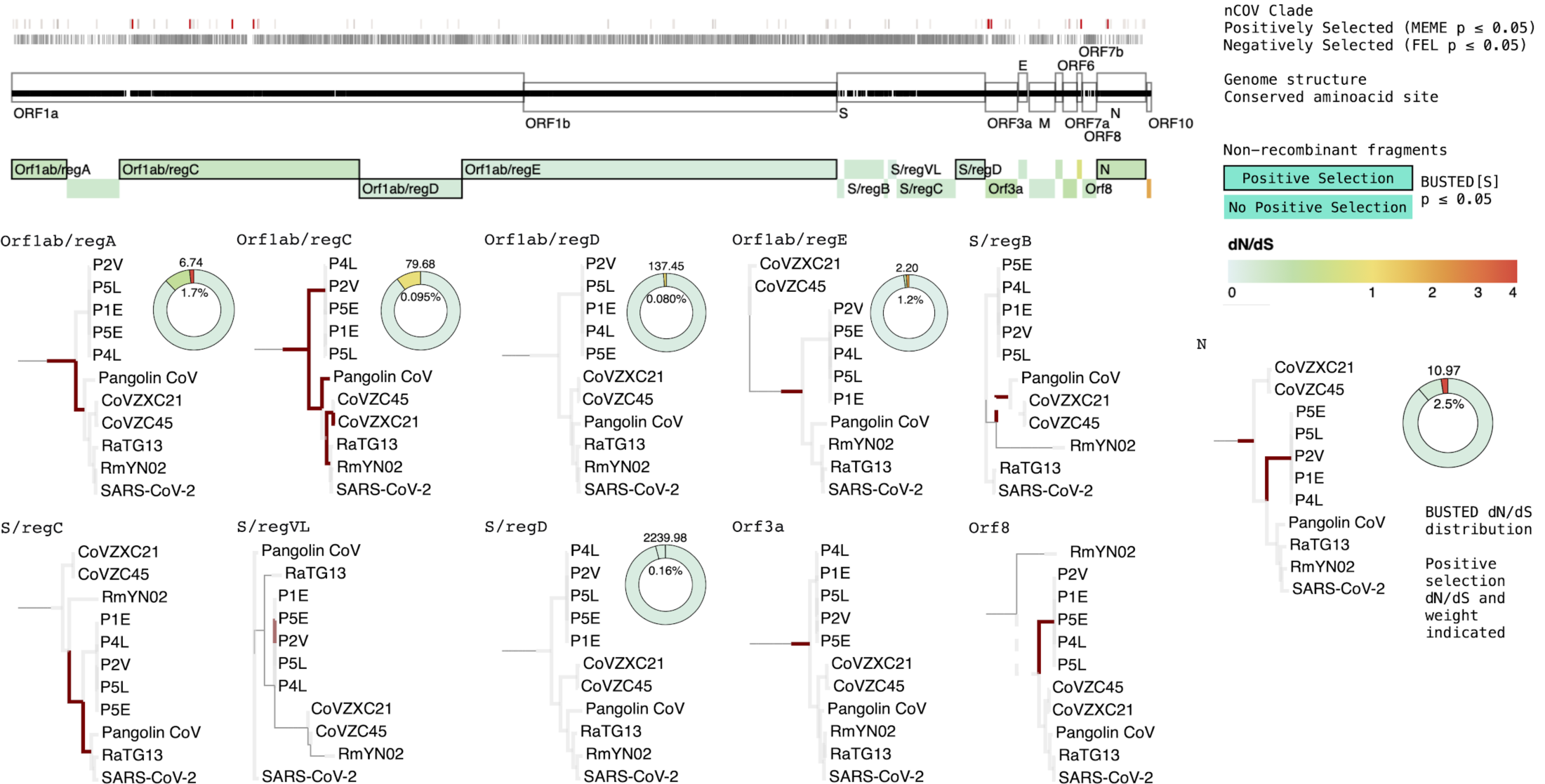
Schematic of the non-recombinant ORF regions used for the nCoV clade selection analyses. Phylogenies with highlighted branches are presented for regions with branch-specific evidence of selection in aBSREL or segment-wide selection using BUSTED[S]. Significance was determined by the likelihood ratio test (p≤0.05) and Benjamini-Hochberg false discovery rate (q≤0.20). Non-nCoV clades are removed for clarity. The inferred dN/dS distribution for each segment found to be under selection using BUSTED[S] is shown as a donut plot (Supplementary Table 4). Three categories of individual sites (conserved, negatively selected, positively selected) are shown as tracks in or above the schematic. For positively selected sites, colouring reflects the fraction of the branches in the nCoV clade inferred to be under selection at the site (gray: smaller, red: larger); further discussion of some of these sites is provided in the text.

To search for branch-specific evidence of selection in the non-recombinant regions, we first used the aBSREL method^36^. Our analysis found that diversifying selection left its imprints primarily in the deepest branches of the nCoV clade or lineage leading to it, with no evidence of selection in the terminal branch leading to SARS-CoV-2 (Figure 2). This is consistent with the non-human progenitor of SARS-CoV-2 requiring little or no novel adaptation to successfully infect humans. Still, no model can detect all signatures of historic genomic adaptation, and mutations which may enable SARS-CoV-2 to infect humans could have arisen by genetic drift in the reservoir host before human exposure.

We next sought evidence of episodic diversifying selection in the nCoV clade using BUSTED[S]^37^, coupled with a hidden Markov model (HMM) with three rate categories to describe site-specific synonymous rate variation (SRV) and allow autocorrelation across neighbouring sites in these rates^38^. Non-recombinant regions of Orf1ab, Spike and ORF N show evidence of episodic diversifying positive selection in the nCoV clade (Figure 2). This finding is consistent with evidence of positive selection operating on Orf1ab in MERS^39^, and Spike and N proteins being essential for antigenic recognition. Eighty-five individual sites were inferred to evolve subject to episodic diversifying selection in the nCoV clade or lineage leading to it (Figure 2; Supplementary Table 3) using the MEME^31^ method. Most of these sites are found on Orf1ab, which is also the longest region. Consistent with BUSTED[S] results, Spike and ORF N have the next greatest number of selected sites (Supplementary Table 3). Interestingly ORF3a has six sites with evidence of positive selection. Little is known about the function of this accessory gene, however, it could be related to immune evasion, like that of other short accessory genes (e.g. Orf3b)^40^. Still, this analysis does not allow attributing specific functional relevance to these individual sites.

The BUSTED[S] method also partitioned synonymous rate variation into three rate classes across the sites. The majority of regions showed large, in some cases more than 20-fold, differences between rate classes, with all three classes representing a substantial proportion of sites for most regions (Supplementary Figure 6), with varying degrees of autocorrelation. This suggests that strong purifying selection is acting on some synonymous sites (e.g., conserved motifs or RNA features), and some synonymous mutations in the SARS-CoV-2 genome may not be selectively neutral or occur at sites that are hypervariable. Some synonymous rate variation may also be attributed to the 5’ and 3’ context-specific mutation rate variation observed in SARS-CoV-2^29^.

### Patterns of CpG depletion in the nCoV clade

Genome composition measures, such as dinucleotide representation and codon usage can also be an informative tool for characterising the host history of a virus^41^. Various host antiviral mechanisms accelerate the depletion of CpG dinucleotides (a cytosine followed by a guanine in the 5’ to 3’ direction) in virus genomes. This is thought to be primarily mediated either through selective pressures by a CpG-targeting mechanism involving the Zinc finger Antiviral Protein (ZAP)^42^ or C to U hypermutation by APOBEC3 cytidine deaminases^43^, and recent work has demonstrated that the CpG binding ZAP protein inhibits SARS-CoV-2 replication in human lung cells^44^. These forces are likely to vary across tissues and between hosts. Thus, a smaller or greater level of CpG depletion in particular viral lineages may be indicative of a switch in the evolutionary environment of that lineage or its ancestors. Although care must be taken to not over-interpret these results^45^.

We examined the CpG representation using the corrected Synonymous Dinucleotide Usage (SDUc) framework, controlling for amino acid abundance and single nucleotide composition bias in the sequences^46^. In Orf1ab, the longest ORF in the genome, all viruses show CpG under-representation (SDUc < 1), while viruses in the nCoV clade have even lower CpG levels than the other *Sarbecoviruses* (Supplementary Figure 7). To further examine this trend, we fitted CpG representation as traits in a phylogenetic comparative approach, using two long putatively non-recombinant regions in the alignment (NRR1 and NRR2; see Methods). Our method identified an adaptive shift favouring CpG suppression in the lineage leading to nCoV clade (Figure 3A). This is coupled with an inflated substitution rate in this lineage compared to the rest of the phylogeny (Figure 3A). We propose that the elevated rate specific to the nCoV clade ancestral lineage can explain some of the variation between different *Coronaviridae* substitution rates, previously attributed to a time-dependent evolutionary rate phenomenon^6^. This shift may indicate a change of evolutionary environment, e.g., host or tissue preference, since CpG depletion following host switches has been observed in other human infecting RNA viruses, such as Influenza B^41^.

**Figure 3.**
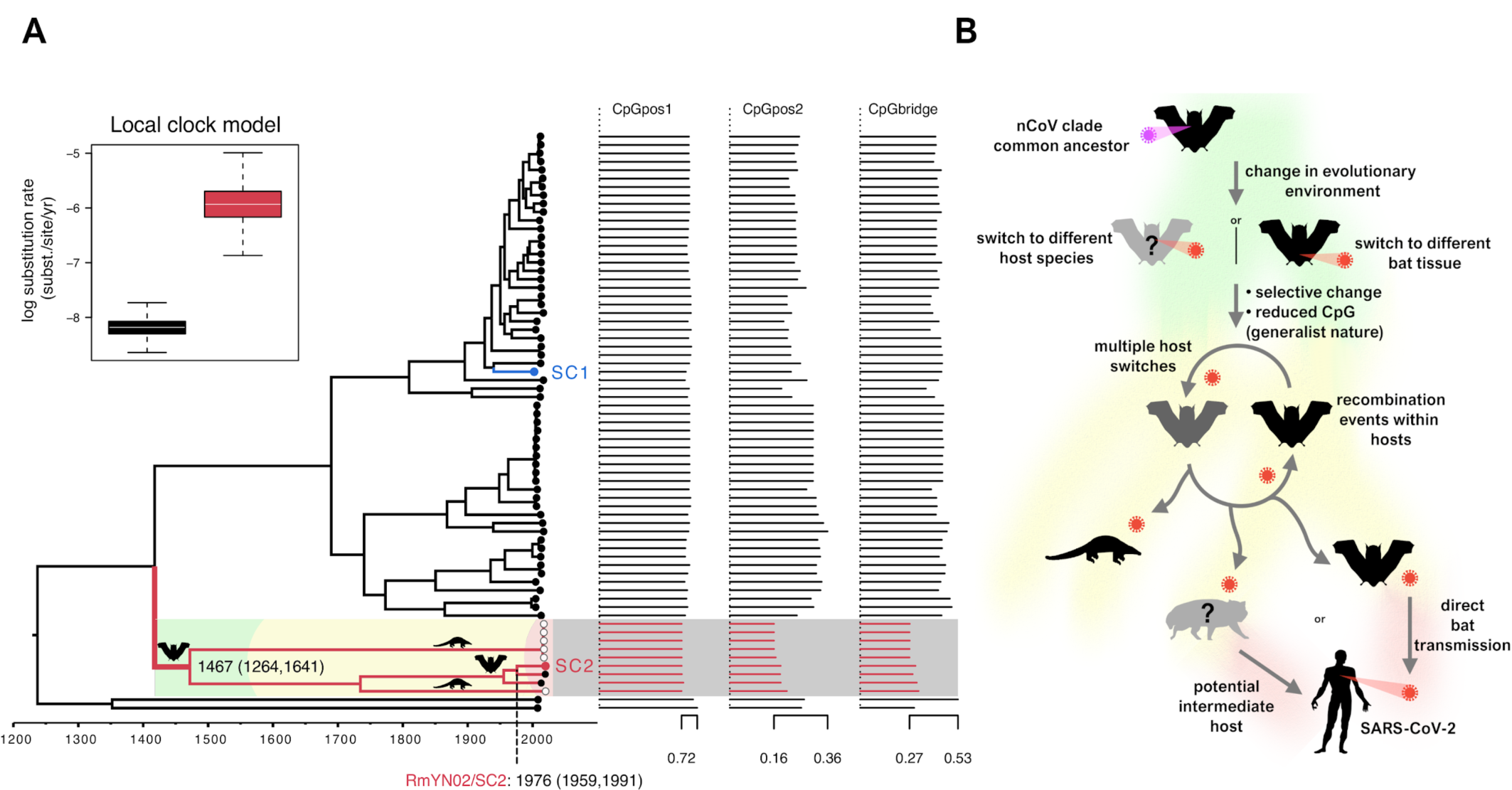
**(A)** Bayesian phylogeny of a modified NRR2 region using different local clocks for the nCoV clade (red branches) and the rest of the phylogeny. Viruses infecting bats are indicated by black circles, pangolins white circles, and SARS-CoV-1 (SC1) and SARS-CoV-2 (SC2) labelled in blue and red, respectively, at the tips of the tree. The inset summarises the substitution rate estimates on a natural log scale for the two-parameter local clock model with colours corresponding to the branches in the tree. The estimated date for the shared common ancestor of the nCoV clade (1467) and the RmYN02/SARS-CoV-2 divergence (1976) are shown with confidence intervals. CpG relative representation for all dinucleotide frame positions (pos1: 1st and 2nd codon positions; pos2: 2nd and 3rd codon positions; bridge: 3rd codon position and 1st position of the next codon) is presented as SDUc values. **(B)** Schematic of our proposed evolutionary history of the nCoV clade and putative events leading to the emergence of SARS-CoV-2.

Zhou et al. (2020)^47^ report a novel bat-infecting *Sarbecovirus* sample, RmYN02, which has the most similar sequence to SARS-CoV-2 of known *Sarbecoviruses* for most of its genome, closer than RaTG13. The part of the RmYN02 Spike ORF that is recombinant falls outside the nCoV clade of the *Sarbecovirus* phylogeny. Molecular dating using BEAST^48^ on a region chosen to be a non-recombinant part of the genome^6^ (Figure 3A, see Methods) indicates that RmYN02 shared an ancestor with SARS-CoV-2 about 1976, strengthening the evidence for a bat to human SARS-CoV-2 emergence. The RmYN02 sequence offers an opportunity to test if the recombination scenario was consistent with the lineage-specific CpG depletion patterns. A sliding window of CpG relative dinucleotide abundance (RDA)^49^ shows that CpG levels of SARS-CoV-2 and RmYN02 only differ at the recombinant region (Figure 4A), further demonstrated by the SDUc values of the nCoV and non-nCoV parts of RmYN02 Spike (Figure 4B). The finding that RmYN02 is a recombinant between the high and low CpG lineages means that viruses from both lineages are co-infecting the same bat species. Both the shift in CpG selection and much of the inference of positive selection at the protein level (Figure 2), coupled with the consistency of the CpG selection pressure throughout the lineage should reflect a single expansion in evolutionary environment in the lineage leading to the nCoV clade. That viruses from this lineage have not subsequently specialised on novel distinct non-bat hosts is also supported by the inability to discover related viruses in wild pre-trafficked malayan pangolins^50^. Together, this suggests the shift at the base of the lineage does not represent specialisation on a host other than bats, but instead an evolved preference towards different host tissues or perhaps circulation in a group of bat species possessing distinct antiviral immunity.

**Figure 4.**
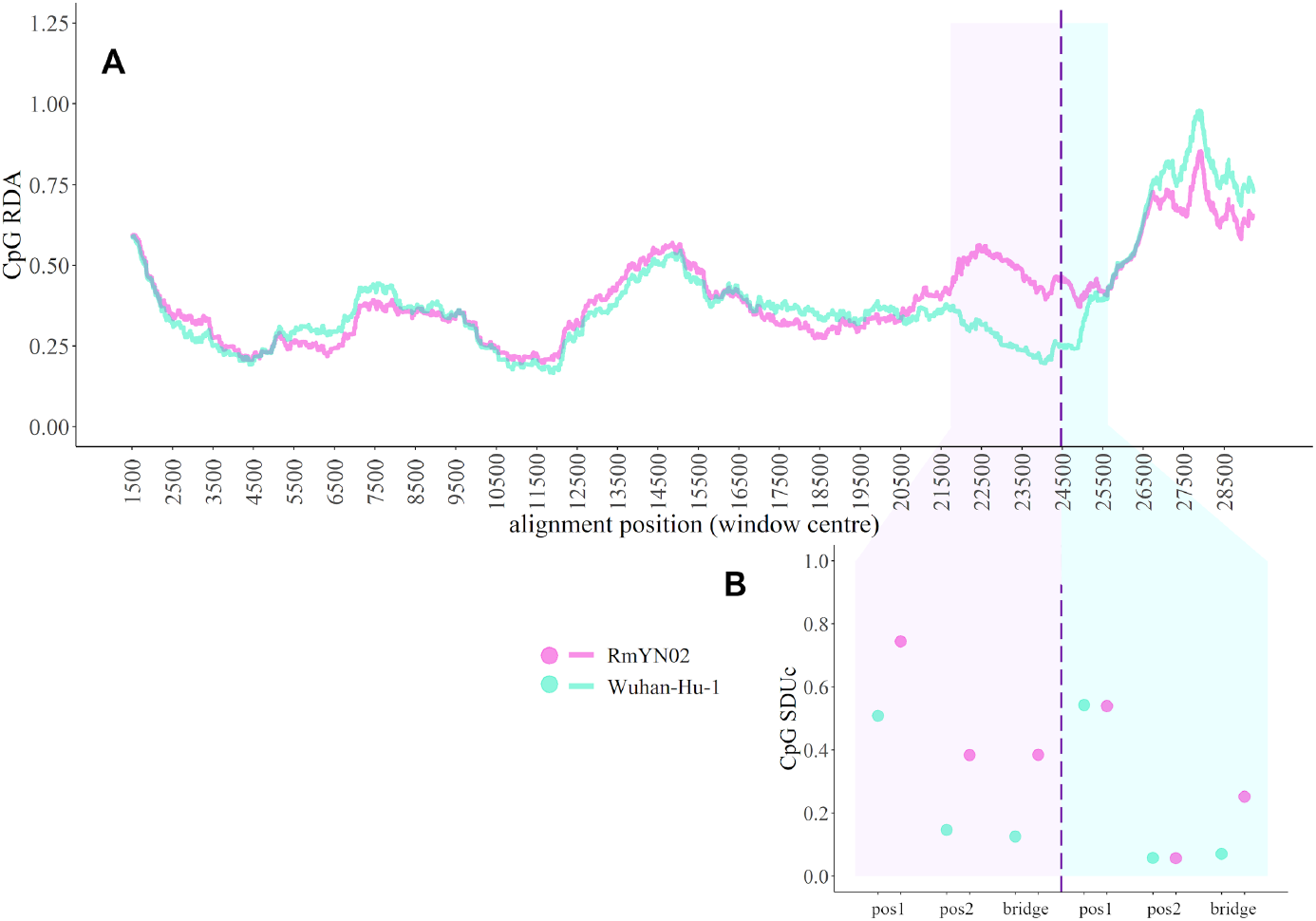
**(A)** 3kb sliding window plot of relative dinucleotide abundance (RDA) across the whole-genome alignment of Wuhan-Hu-1 (turquoise) and RmYN02 (magenta). Shaded regions depict the Spike ORF region in the alignment. The dashed line indicates the inferred RmYN02 Spike recombination breakpoint, splitting the shaded region into non-nCoV (pink) and nCoV (blue). **(B)** SDUc values calculated for each frame position of the two RmYN02 Spike non-recombinant regions and the corresponding Wuhan-Hu-1 regions.

## Conclusion

The evidence of positive selection on the nCoV clade SARS-CoV-2 emerged from – coupled with the adaptive shift in CpG composition in this lineage and lack of evidence of ramped up diversifying selection in human circulating SARS-CoV-2 – indicates that significant events in SARS-CoV-2 evolution occurred prior to spillover into humans. Though we do find evidence of some ongoing positive selection and minor optimisation to the human population (Figure 1E), the small number of sites involved and lack of selective sweeps since sequencing began demonstrates the virus was already a highly capable human pathogen. Changes specific to SARS-CoV-2 that are significant for the virus’ success in humans include: (i) the specific amino acid replacements observed in receptor binding domain (RBD) in Spike. However, their shared ancestry with pangolin viruses is due to complex recombination events involving bat viruses^6^ indicating that this RBD-phenotype predates any modern contact between bats and humans and so is not associated with the recent emergence, and (ii) the acquisition of a polybasic cleavage site in Spike due to non-homologous recombination involving bat viruses outside of the nCoV lineage (Supplementary Text 4, Supplementary Figure 10). Note, we cannot discount this latter event occurred in either a co-infected intermediate species or even a co-infected human. Recombination between a proximal ancestor of RmYN02 and a non-nCoV *Sarbecovirus* further indicates the viruses in this subgenus co-circulate in the same bat reservoirs. The apparent ‘success’ of these bat viruses in several species following cross-species transmission, without significant genomic changes, supports the hypothesis that the SARS-CoV-2 progenitor is from a viral lineage with a relatively generalist nature (Figure 3B). We suggest that the early ancestors of the nCoV clade – existing hundreds of years ago (Figure 3A, Supplementary Table 5) – developed this generalist phenotype through an expansion in their evolutionary environment (host switch, or tissue tropism) within bat species, allowing for efficient spillover events. Given there remain large gaps in our knowledge of SARS-CoV-2’s recent non-human origins, as the closest bat viruses are relatively divergent in time^6^, the generalist nature of nCoV clade members dramatically increases the probability of there being wild mammals we have yet to sample infected with nCoV-like viruses. Serological studies of communities in China that come into contact with bats indicate that incidental and ‘dead-end’ spillover of SARS-like viruses into humans do occur^51,52^. Due to the high diversity and generalist nature of these *Sarbecoviruses*, a future emergence, or recombination event is likely, and such a ‘SARS-CoV-3 virus’ could be sufficiently divergent to evade either natural or vaccine-acquired immunity, as demonstrated for SARS-CoV-1 versus SARS-CoV-2^53^. We must therefore dramatically ramp up surveillance at the human-animal interface, monitor carefully for both reverse-zoonosis establishing SARS-CoV-2 in new animal reservoirs and future SARS-CoV emergence in the human population. SARS/COVID-19 causing viruses are here for the long-term.

## Methods

### SARS-CoV-2 GISAID sequence filtering

To reduce the impact of sequencing errors on selection analysis the data from GISAID was filtered by excluding all sequences which meet any of the following criteria: any sequence of length less than 29000 nucleotides; any sequence with a non-human host (e.g., bat, pangolin); any sequences with ambiguous nucleotides in excess of 0.5% of the genome; any sequences with greater than 1% divergence from the longest sampled sequence (Wuhan-Hu-1); and any sequence with stop codons. Unique (identical sequences are collapsed into a single group) codon-aligned sequences are translated to amino-acids and aligned with MAFFT^54^. The amino-acid alignment is mapped back to the constituent nucleotide sequences. Only unique haplotypes are retained for comparative phylogenetic analyses, since including identical copies is not informative for these types of inference.

### SARS-CoV-2 positive selection

We initially utilised methodology which incorporated the full phylogeny and found misleading signatures such as sequencing errors and lab-based recombination on the terminal branches, which confounded the software (Supplementary Text 2-3). These errors are reflected in the elevated dN/dS ratio observed on the terminal branches (Figure 1). We subsequently used the software MEME^31^ to infer positive selection, using an implementation which only considered internal branches to infer positive selection. MEME uses a maximum likelihood methodology and performs a likelihood ratio test for positive selection on each site, comparing modes which allow or disallow positive diversifying selection at a subset of branches (dN/dS>1).

### *Sarbecoviruses* alignment and recombination

To avoid the confounding effects of recombination, we have analysed each open reading frame (ORF) separately and divided the Orf1ab and Spike ORFs into putative non-recombinant regions, based on the seven major recombination breakpoints presented in Boni et al. (2020)^6^. This produces five non-recombinant regions for Orf1ab (regions A to E) and five regions for Spike (regions A to D, and the variable loop - region VL). The protein sequences of the non-recombinant regions SARS-CoV-2, SARS-CoV-1 and 67 closely related viruses with non-human hosts (bats and pangolins; Supplementary Table 6) were aligned using MAFFT version 7 (L-INS-i)^54^. Subsequent manual corrections were made on the protein alignments and pal2nal^53^ was used to convert them to codon alignments. Phylogenies for each codon alignment were inferred using RAxML with a GTR+Γ nucleotide substitution model^55^.

### *Sarbecovirus* selection analysis

We used an array of selection detection methods to examine whether the lineage leading to SARS-CoV-2 has experienced episodes of diversifying positive selection. Each non-recombinant region was examined separately. We separated each region’s phylogeny into an nCoV and non-nCoV/SARS-CoV-1 lineage. The nCoV clade includes SARS-CoV-2 and the viruses that it is phylogenetically most closely related to. These are the bat-infecting CoVZC45, CoVZXC21, RmYN02 and RaTG13, and the pangolin-infecting Pangolin-CoV and P2V, P5L, P1E, P5E, P4L cluster. Note, some recombinant regions of the first three viruses do not belong to the nCoV clade.

We tested for evidence of episodic diversifying selection on the internal branches of the nCoV clade using BUSTED[S], accounting for synonymous rate variation (SRV) as described in Wisotsky et al. (2020)^37^. We developed an extension to BUSTED[S], that included a hidden Markov model (HMM) with three rate categories to describe site-specific synonymous rate variation (SRV)^38^. This HMM allows explicit incorporation of autocorrelation in synonymous rates across codons. This autocorrelation would be expected if selection or mutation rate variation were spatially localised within ORFs. The rate switching parameter between adjacent codons of the HMM describes the extent of autocorrelation, with values under 1/N (N = number of rate classes) suggestive of autocorrelation. Standard HMM techniques (e.g. the Viterbi path) applied to these models can reveal where the switches between different rate types occur, thereby partitioning the sequence into regions of weaker or stronger constraint on synonymous substitutions.

The aBSREL method^36^ was used on all branches of the nCoV clade to determine which specific branches drive the inference of selection. Finally, we examined which specific codon sites are under negative selection on average over the nCoV clade using FEL^25^, and under pervasive or episodic diversifying positive selection on the nCoV clade using MEME^31^. P-values of ≤0.05 for the likelihood ratio tests, specific to each method, were taken as evidence of statistical significance. All selection analyses were performed in the HyPhy software package v.2.5.14^56^.

### CpG depletion

To quantify over/under representation of CpG dinucleotides in the *Sarbecovirus* genomes we developed a modified version of the Synonymous Dinucleotide Usage (SDU) metric^46^, which now accounts for biased base composition. The original SDU metric compares the observed proportion of synonymous CpG, *o*, for each pair of frame positions, *h*, in a coding sequence to that expected under equal synonymous codon usage, *e*, for each amino acid (or amino acid pair), *i*, that can have CpG containing codons (or codon pairs). The SDU metric is the mean of these ratios weighted by the number of informative amino acids (or pairs), *n*, in the sequence (Equation 1).

To incorporate the biased, and variable base composition of SARS-CoV-2 and other *Sarbecoviruses*^29^, here we have estimated expected codon usage based on each virus’s whole-genome nucleotide composition. We term this new metric the corrected Synonymous Dinucleotide Usage (SDUc). We use observed base frequencies from each virus to generate the corrected null expectation of the metric, *e’*, instead of assuming equal usage, (Equation 1). The expected proportion, *e’*, for every amino acid / amino acid pair was estimated by randomly simulating codons based on the whole-genome single nucleotide proportions of each virus. This *e’* was then used for all SDUc calculations of the corresponding virus.

As this metric is susceptible to error when used for short coding sequences, we applied SDUc on the longest ORF, Orf1ab, of all the viruses. To estimate the extent of phylogenetic independence between synonymous sites across SDUc datapoints, we measured the pairwise synonymous divergence (Ks) between viruses. Pairwise Ks values were calculated using the seqinr R package^57^ which utilises the codon model of Li (1993)^58^, demonstrating the partial but not complete independence within the two lineages. The Ks median and maximum is 0.54 and 0.89 within the nCoV clade, and 0.34 and 1.09 respectively within the non-nCoV/SARS-CoV-1 clade.

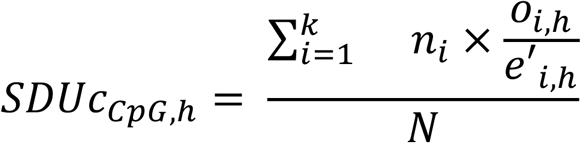

(Equation 1; N = total number of informative amino acids)

### Spike recombination analysis

To determine the recombination breakpoint on the Spike ORF of the RmYN02 virus we used the RDP5 method suite^59^, implementing seven methods: RDP, GENECONV, Chimaera, MaxChi, BootScan, SiScan and 3seq. We first performed the analysis on the whole-genome alignment of the *Sarbecoviruses* and then determined the relevant breakpoint within the Spike ORF by rerunning the method on the Spike-only alignment. The accepted breakpoint (position 24058 in the RmYN02 genome) was consistently called by six out of the seven tested methods (RDP, GENECONV, Maxchi, Chimaeara, SiSscan, 3seq). Similarly, the preceding breakpoint of the non-nCoV region was called at position 21248 of the RmYN02 genome (before the start of the Spike ORF).

### Bayesian inference under a local molecular clock

To assess the overall non-recombinant phylogenetic relation between the viruses, we used the longest non-recombinant genomic regions described in Boni et al. (2020)^6^, NRR1 and NRR2, including RmYN02 in the alignments. Based on the recombination breakpoints determined above for RmYN02, NRR2 was adjusted to end at position 21266 of Wuhan-Hu-1 - instead of 21753-corresponding to the start of the RmYN02 recombinant region (RmYN02 position 21248). Time-measured evolutionary histories for NRR1 and NRR2 were inferred using a Bayesian approach, implemented through the Markov chain Monte Carlo (MCMC) framework available in BEAST 1.10^48^. Motivated by the observation of a higher root-to-tip divergence for the nCoV clade (Supplementary Figure 8) and by the CpG trait shift on the branch ancestral to the nCoV clade (see below), we specified a fixed local clock^60^ that allows for a different rate on the branch leading to the nCoV clade. In the absence of strong temporal signal, we specified an informative normal prior distribution (with mean = 0.0005 and standard deviation = 0.0002) on the rate on all other branches based on recent estimates under a relaxed molecular clock^61^. To maintain identifiability under the local clock model, we constrained a Kenyan (KY352407) bat virus as outgroup for the viruses from China in NRR1, and the Kenyan as well as a Bulgarian bat virus (NC_014470) as outgroup for the viruses from China in NRR2. We partitioned the coding regions of NRR1 and NNR2 by codon position and specified an independent general time-reversible (GTR) substitution model with gamma-distribution rate variation among sites for each of the three partitions. We used a constant size coalescent model as tree prior and specified a lognormal prior with mean = 6.0 and standard deviation = 0.5 on the population size. Three independent MCMC analyses were run for 250 million states for each data set. We used the BEAGLE library v3^62^ to increase computational performance. Continuous parameters were summarized and effective sample sizes were estimated using Tracer^63^. Divergence time estimates for the nCoV clade are summarised in Supplementary Table 5. Trees were summarized as maximum clade credibility (MCC) trees using TreeAnnotator and visualized using FigTree.

### Identifying shifts in CpG content

Shifts in CpG content were identified using a phylogenetic comparative method that infers adaptive shifts in multivariate correlated traits^64^. This approach models trait evolution on phylogenies using an Ornstein-Uhlenbeck (OU) process and uses a computationally tractable version of the full multivariate OU model (scalar OU) for multivariate traits. Estimates of the shift positions are obtained using an Expectation-Maximization (EM) algorithm. The shift positions are estimated for various numbers of unknown shifts and a lasso-regression model selection procedure identifies the optimal number of shifts. We applied the procedure to the MMC trees for NRR1 and NRR2 with their respective CpG SDUc values (ln transformed as required by the phylogenetic approach). The trees were forced to be ultrametric, as required for the scalar OU model, by extending all the external edges of the trees to match the most recently sampled tip. Using this procedure, we identified three and two CpG shifts in both NRR1 and NRR2 respectively (Supplementary Figure 9), with the shift on the branch ancestral to the nCoV clade being the only consistent one identified in both genomic regions (Figure 3A).

## Acknowledgements

We would like to thank all the authors who have kindly deposited and shared genome data on GISAID. Credit also needs to be given to the surveillance projects for generating the genome data that is available in GenBank and to the software developers for making the tools we’ve used freely available. A table with genome sequence acknowledgments can be found in the supplementary material. We thank Alex Gunnarsson, Xiaowei Jiang, Joseph Hughes and Kyriaki Nomikou for thankful comments on the manuscript. DR and SL are funded by the MRC (MC_UU_1201412). SLKP and SW are supported in part by the NIH (R01 AI134384 (NIH/NIAID)) and the NSF (award 2027196). PL acknowledges funding from the European Research Council under the European Union’s Horizon 2020 research and innovation programme (grant agreement no. 725422-ReservoirDOCS), the European Union’s Horizon 2020 project MOOD (874850), the Wellcome Trust through project 206298/Z/17/Z (The Artic Network) and the Research Foundation -- Flanders (’Fonds voor Wetenschappelijk Onderzoek -- Vlaanderen’, G066215N, G0D5117N and G0B9317N).

