## Supplementary text, tables 2 and 5, and figures 1 to 10 for "Natural selection in the evolution of SARS-CoV-2 in bats, not humans, created a highly capable human pathogen"

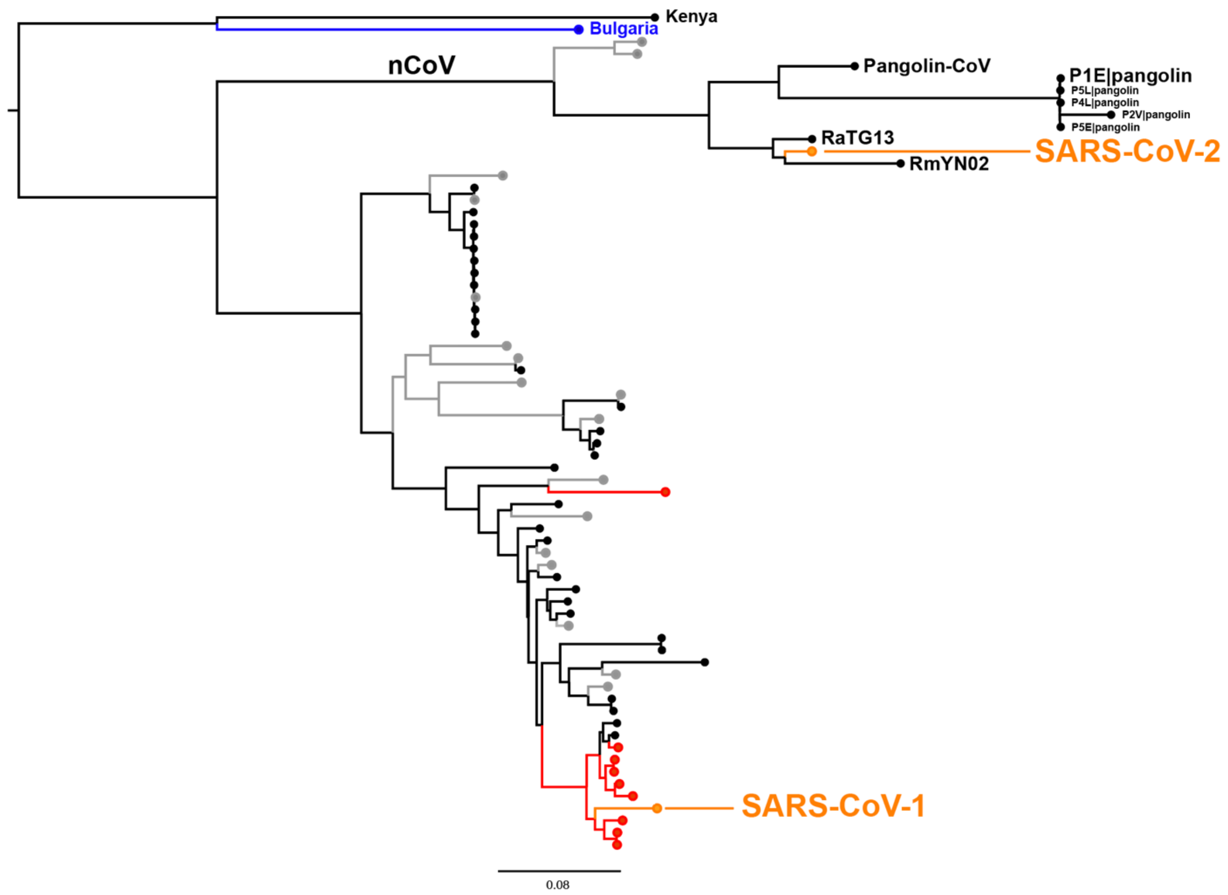


***Supplementary Figure 1***. *Phylogenetic tree (from RAxML*[1] and visualized with FigTree, http://tree.bio.ed.ac.uk/software/figtree/*) showing the relationship of SARS-CoV-1 & -2 (orange) to related bat and pangolin Sarbecoviruses* using whole genome alignments. *Grey, red and blue variants are coloured according to Letko et al. (2020)*[2] *who showed experimentally some viruses are able to use human ACE2 (red), while some require exogenous protease treatment in vitro (grey); the red outlier is a known recombinant. Black indicates not tested by Letko et al, 2020, while the virus in blue, sampled in Bulgaria, could not be induced to infect human cells. The scale bar is in expected nucleotide substitutions per site.*

### Supplementary Text 1: Inferring population demography from distribution of allele frequencies

#### Fitting exponential and constant population size expectations to real data

Population expansion is known to increase the relative proportion of mutations at low frequency. Following our observation that most variation in SARS-CoV-2 data sets is at very low frequency, we compared the observed distribution of SNP frequencies to expectations under two demographic models. An exponential growth model fits much better than a stable population size model, consistent with the recent spread of the virus across the globe. The fit appears to be better than that produced by Lythgoe *et al*. from within-host data[3]. This probably reflects that within-host data is more prone to sequencing errors than consensus base calls especially at low frequencies, or perhaps reflects a super-exponential growth rate within hosts that violates the expectation given by the 1/x^2 equation (personal communication Luca Ferretti).

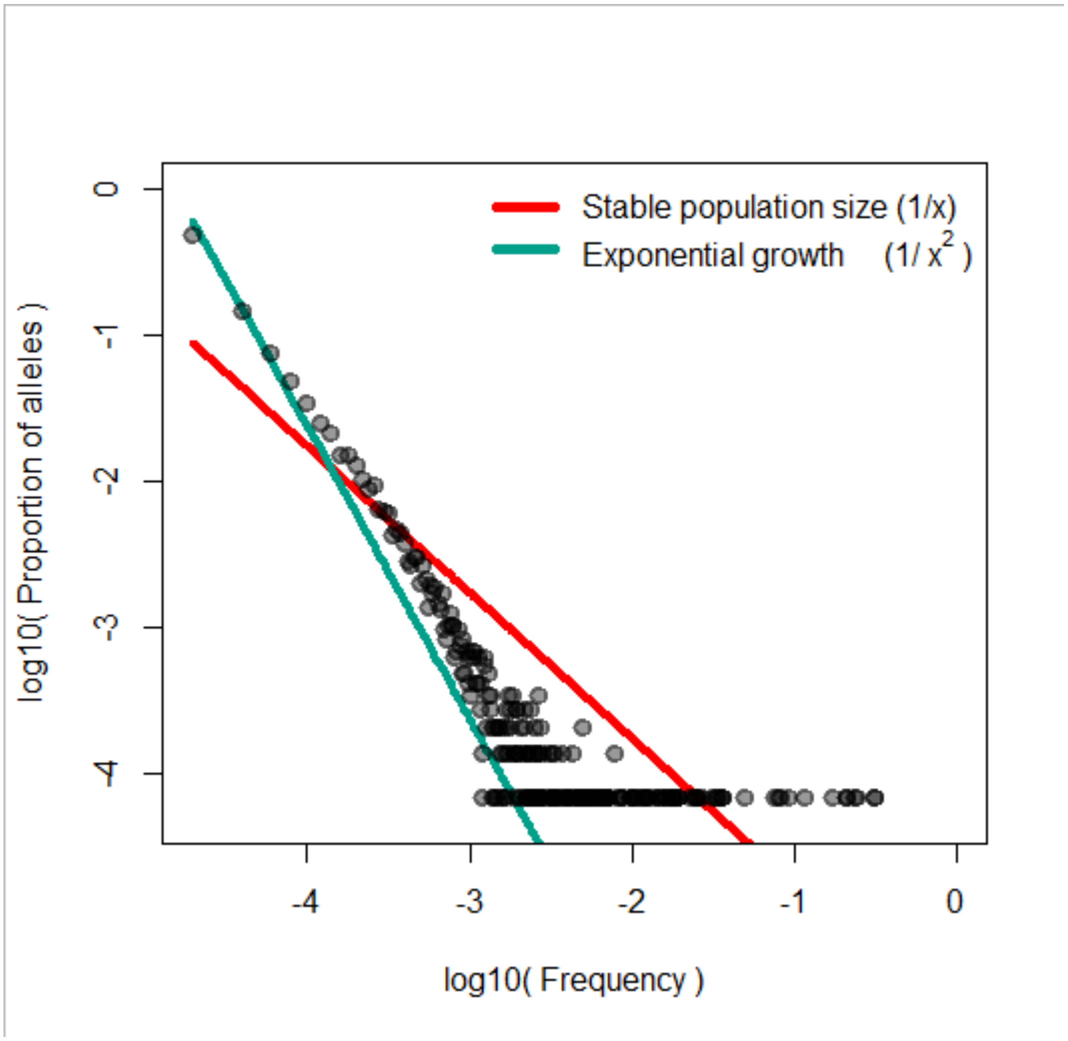

***Supplementary Figure 2.*** Log-log plot of the proportion of minor alleles at each frequency, compared with expectations under two demographic models[3,4]. Allele frequencies were calculated across all ORFs, with each ORF separately passed through QC filters (see main methods) exact number of sequences varies across genomic regions, expectations were calculated assuming 48954 sequences, representing the median number across ORFs (data retrieved from GISAID June 28th, 2020).

### Supplementary Text 2. Signals of positive selection in SARS-CoV-2

**Searching for positive selection within the SARS-CoV-2 outbreak radiation.** We initially used the Bayesian FUBAR software[5] from the HyPhy package to identify sites exhibiting signatures of diversifying positive selection in the SARS-CoV-2 outbreak data. Such signatures are suggestive of the virus undergoing adaptation to humans in the pandemic. FUBAR detects positive selection by looking for codons which have an elevated rate of nonsynonymous protein coding substitutions (dN) relative to synonymous substitutions (dS). It allows both synonymous and non-synonymous rate variation across codons, as has been observed in SARS-CoV-2 evolution[6], but assumes that selective pressures are constant through time, and across all branches (allowing no branch-to-branch variation). It estimates a posterior probability that each site is under positive selection across the phylogeny (dN/dS >1), with a posterior probability >0.9 used as the threshold for significance, as suggested by the authors[5]. The input data for this software was a concatenated coding alignment of 396 sequences from GISAID up to March 16$^{th}$ 2020 (Supplementary Table 1) and using a tree generated in RAxML[1] under the GTR+Γ model. This should be a sufficient number of variants to capture the emergence of SARS-CoV-2 and any early associated adaptations. This analysis reported ten sites as showing significant evidence of positive selection across the pandemic phylogeny (Supplementary Table 2). Due to the low diversity in these 396 SARS-CoV-2 samples, there is limited power to confidently estimate the synonymous and nonsynonymous substitution rate for each codon. This means that statistical power to identify positive selection in the form of dN/dS>1 for any given codon is limited, and the posterior distribution should be flat. The presence of statistically significantly signatures of positive selection is therefore somewhat surprising.

**Supplementary Table 2**. The location and posterior probabilities of the ten mutations detected by the FUBAR selection analysis.

| Codon in concatenated alignment | ORF | mutation | Posterior probability |
|---|---|---|---|
| 476 | NSP2 | I296V | 0.935871674 |
| 1599 | NSP3 | L781F | 0.943645748 |
| 3606 | NSP6 | L37F | 0.975858139 |
| 7461 | Spike | V367F | 0.952007026 |
| 7708 | Spike | D614G | 0.939001013 |
| 7954 | Spike | V860Q | 0.919218684 |
| 7955 | Spike | L861K | 0.944083041 |
| 8720 | ORF-M | D3G | 0.938851732 |
| 9248 | ORF-8 | L84S | 0.939347212 |
| 9577 | ORF-N | I292T | 0.952894581 |

As recombination is known to confound selection analyses such as the methods in the HyPhy package[7], the maximum likelihood recombination detection software GARD[8] was used to test for recombination before performing selection analysis. This software searches for recombination by introducing potential breakpoints and optimising tree topologies either side of the new breakpoint. If the Akaike information criterion (AIC)[9] is improved by the optimisations with breakpoints in, this provides significant evidence of recombination. If significant evidence of recombination is found, the method can then generate multiple non-recombinant partitions in the sequence alignment for use in downstream analyses. However, if the samples are highly related, as in the SARS-CoV-2 dataset, this phylogeny-based approach is limited in power as each recombination event introduces a large number of additional number of parameters, substantially penalising the AIC[9]. To detect recombination with more power for closely related samples, we also used the pairwise homoplasy index[10], which tests for excessive homoplasies. However, this method cannot tell if homoplasies are due to recombination or convergent evolution through parallel adaptation due to shared selection pressures.

To understand the specific mutational patterns that might explain these significant results, we looked at where in the phylogeny these putatively positively selected mutations were occurring. For all but two of the ten positive selected codons (Spike codons 860 and 861 highlighted in red in Supplementary Table 2), this signal was being driven by apparent convergent evolution (or homoplasy) in the tree, with the same mutation occurring in parallel across the phylogeny. To investigate whether this observation was truly due to independent events or because of recombination signatures in the SARS-CoV-2 outbreak tree, we firstly determined if the samples with these convergent mutations were geographically correlated. As selective pressure acting on an untreatable novel zoonotic virus is likely to be globally shared (adaptation to humans), but recombination requires co-localisation of viruses in the time and space, geographic clustering would be a good indication that these mutations are not independent.

The homoplasies driving ORF8 L84S and ORF1ab L1599F mutations were both found in South Korean isolates, and each of the two instances of ORF N I292T were found in the Netherlands. This geographic clustering was suggestive of recombination and was investigated further, see below.

**Recombination or selection**. For the two FUBAR-flagged sites, Spike codons 860 and 861, that did not show any homoplasies, both signals could be attributed to the same run of four neighbouring U to A mutations spanning the two codons. These mutations were found in only a single sample: EPI_ISL_408485 from Beijing and have not been observed since (to date 8/5/2020). This suggests that they were either sequencing errors or a large single mutation spanning two codons, which has not subsequently spread. Multiple nucleotide changes within a single codon should be rare and sequencing error is a plausible explanation.

The positive selection signature at NSP6 codon 37 can be explained by multiple homoplasies of G to U mutations at nucleotide 11083. This mutation is found in four distinct haplotypes (Supplementary Figure 1) across different areas of the phylogeny. There are flanking mutations on both sides of this site shared by sequences which both possess and do not possess the 11083 mutation. For this to occur under a recombination scenario, multiple breakpoints would be required for each homoplasy. These observations therefore are not most parsimoniously explained by recombination alone. The presence of this mutation across the tree could be driven by either positive selection, parallel sequencing error or hypermutability through polymerase slippage.

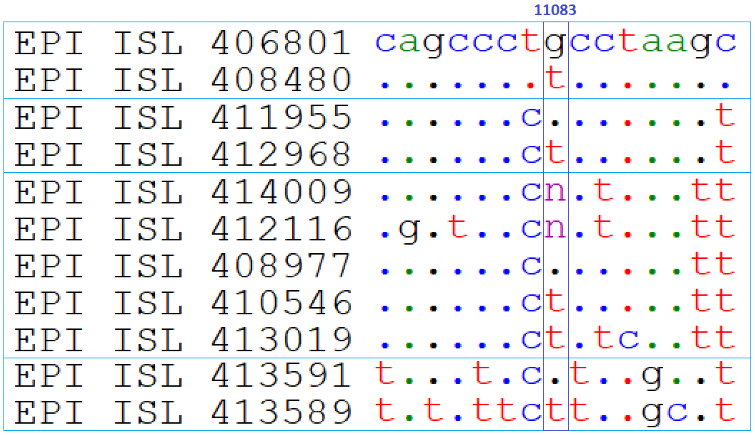


***Supplementary Figure 3.*** *Alignment of variable sites (removing invariant sites across these samples) surrounding the ORF1A L3606F recurrent G->T mutation. It appears to independently originate on four genetic backgrounds. The presence of 'n's in a few of the samples suggests that sequencing ambiguity is common in this region.*

Both the ORF8 codon 84 and ORF1ab codon 1599 positive selection signals appear to be raised by a single South Korean sample (GISAID accession 413017). This sample possesses two derived mutations either side of a hypothesised breakpoint. These pairs of derived mutations belong to samples with different haplotypes (Supplementary Figure 2). Therefore the 413017 sample appears to be a recombinant between sample 413018 and 413513 or 412871. As both 413017 and 413018 were sequenced by the same laboratory and released at the same time, this recombination event may be an artefactual product of laboratory cross-contamination.

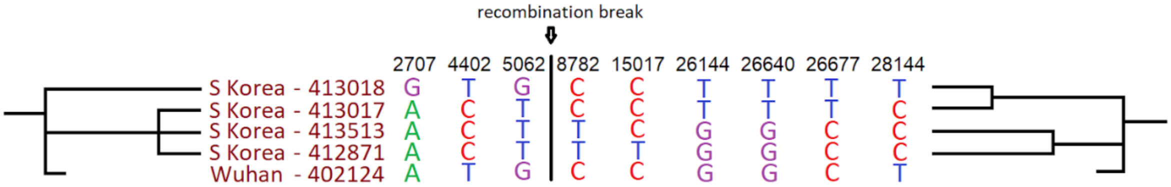


***Supplementary Figure 4.*** *Alignment of variable sites in the whole genome alignment, with base positions shown above. The putative recombinant South Korean sample 413017 shows mismatching topologies across its genome, clustering with different South Korean samples either side of the inferred breakpoint. The Wuhan sample 402124 was collected on 30/12/2019, and shows no unique mutations, it should thus represent the ancestral state of the four chosen samples.*

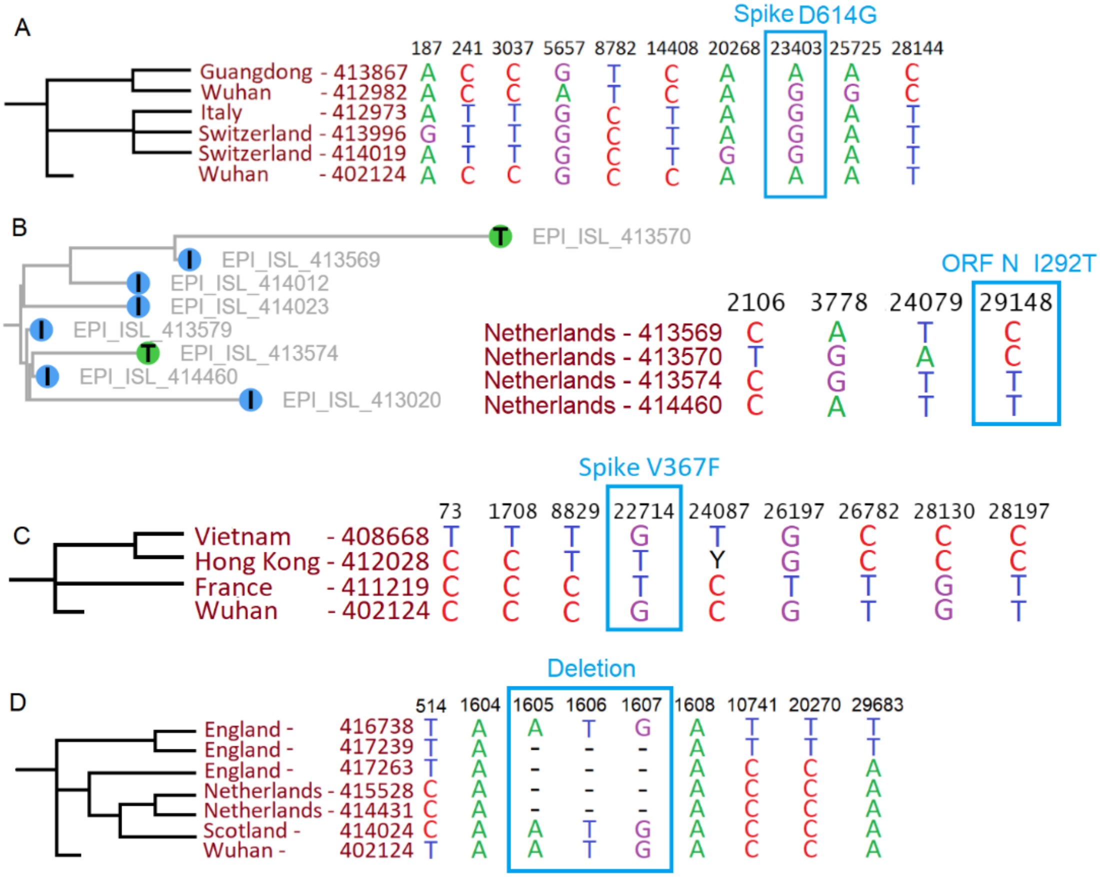


***Supplementary Figure 5.*** *A) Spike D614G replacement: sites either side of D614G show derived mutations in Wuhan 412982 congruent with the tree, suggesting that it is not a recombinant, or has multiple breakpoints. B) N ORF I292T: both Netherlands samples with the mutation were sequenced by the same Dutch laboratory and released at the same time. C) Spike V367F homoplasy at a single site in Hong Kong, unlikely to be a recombinant. D) An observed insertion homoplasy in newer data.*

The Spike D614G signal was driven by apparent convergent evolution between one Wuhan sample (412982) in addition to the main lineage containing 86 samples. This sample shares mutations with its closest related sequence (Guangdong 413867) on both sides of this homoplasy (Supplementary Figure 5A), suggesting it is not the result of recombination. No newly sequenced samples uploaded up to 27/4/2020 containing the D614G mutation clustered with the Wuhan 412982 sample, suggesting that this haplotype did not spread or that this homoplasy is driven by sequencing error. Additional sequences displaying apparent convergent evolution at this site have since been sequenced, these have been taken as evidence of positive selection[11]. However, given that this mutation now occurs in 59% of sequenced samples (as of 27/4/20), it will be one of the mutations most likely to be variable if multiple viral genotypes are present following laboratory contamination or in mixed infections, and so most prone to being shuttled onto new backgrounds by recombination. Therefore, whilst high frequency mutations are the most important to study, they are also the most prone to misleading homoplasies, and must be analysed with the most caution.

The N ORF 292 site detected by FUBAR is driven by a similar convergent evolution event history. However, both samples exhibiting the same derived I to T mutation (Supplementary Figure 5B; GISAID IDs 413570 and 413574) were sequenced by the same Dutch laboratory and released at the same time, again suggesting that laboratory cross contamination is a likely driver. However, unlike South Korean sample 413017, there is only one shared derived mutation (codon 292), and therefore the genomic evidence for recombination in these samples is weaker.

The ORF M D3G mutation was found in three samples, 414010, 414017, and 413999 from England, Scotland, and Switzerland respectively. The positive selection signal was driven by sample 414010 from England, which exhibited a homoplasy at ORF N codon 156, representing the A156S mutation, which is shared with two samples from the Netherlands 414450 and 414457. More recent samples since this analysis which exhibit both the ORF M D3G and ORF N A156S mutation have been sequenced in England (e.g. 449635) suggesting that this sample represents a true recombination/ convergent evolution event which was transmitted. Which of these two codons represents the convergent mutation/ recombination event is unclear due to the low levels of divergence between lineages at the early stage of the pandemic when this event occurred. Additionally, samples exhibiting only one of ORF M D3G or N A156S, but not both, are observed in more recent sequencing data, e.g. English sample 461979 for the former and Indian sample 475029 for the latter.

The Spike V367F replacement signal was driven by apparent convergent evolution between four French samples sequenced in January and a Hong Kong sample 412028, which shows shared variation either side of the homoplasy suggesting it is not a recombinant (Supplementary Figure 5C). Looking through more recent data shows additional homoplasies in a neighbour joining tree. Additionally, newly generated sequences since the FUBAR analysis cluster around the Hong Kong sample, further suggesting it is not a laboratory generated sequencing error. This site was also flagged in our updated methodology in mid-May (main text Figure 2E), however this substitution has not been seen for months, suggesting that it stochastically been lost. It might be speculated that the multiple origins of this amino acid replacement might be driven by positive selection within hosts, and its loss may have been driven by negative selection between hosts. Trade-offs of this kind have been observed in HIV-1[12].

Subsequent scans of newer data have revealed additional evidence of laboratory recombination events (see Supplementary Figure 5D). Given these observed issues with the data, it is clear that analysis for positive selection should not consider terminal branches in searching for dN/dS>1, as these are prone to sequencing error artefacts, and analyses should instead only utilise internal branches.

### Supplementary Text 3. Frequency-based analysis of SARS-CoV-2 polymorphisms

In addition to searching for positive selection, we investigated if signatures of purifying selection on segregating variation in the current SARS-CoV-2 data could be observed (sequences as of 14/5/20). We compared the relative frequencies of nonsynonymous and synonymous mutations in the pandemic data. Codons with multiple mutations present were discarded from the analysis to avoid ambiguity in the order of mutations and simplify synonymous/nonsynonymous classification.

Most mutations of both classes are at very low frequency (main text Figure 2), indicative of the viral population expansion that the pandemic has undergone. dN/dS was approximately 0.6 in singletons, suggesting that 40% of nonsynonymous mutations are strongly deleterious and therefore never observed in the population. There is a weak observable trend towards a higher proportion of mutations being synonymous at the highest frequency intervals, suggestive of some ongoing selection against circulating amino acid replacements in the pandemic. This observation may be partially driven by sequencing errors which are not transmitted and so are at low frequency. These sequencing errors are likely to have a dN/dS value of 1, which may make the estimate that 40% amino acid replacements are strongly deleterious an underestimate of the true value. However, the decline in nonsynonymous/synonymous ratio occurs across the range of frequencies, suggesting that sequencing errors alone are not driving the trend. It is important to consider that the observed frequencies are likely to differ from true global frequencies due to biased sampling of infections in the pandemic[13], and so we caution against overinterpretation of specific mutation frequencies.

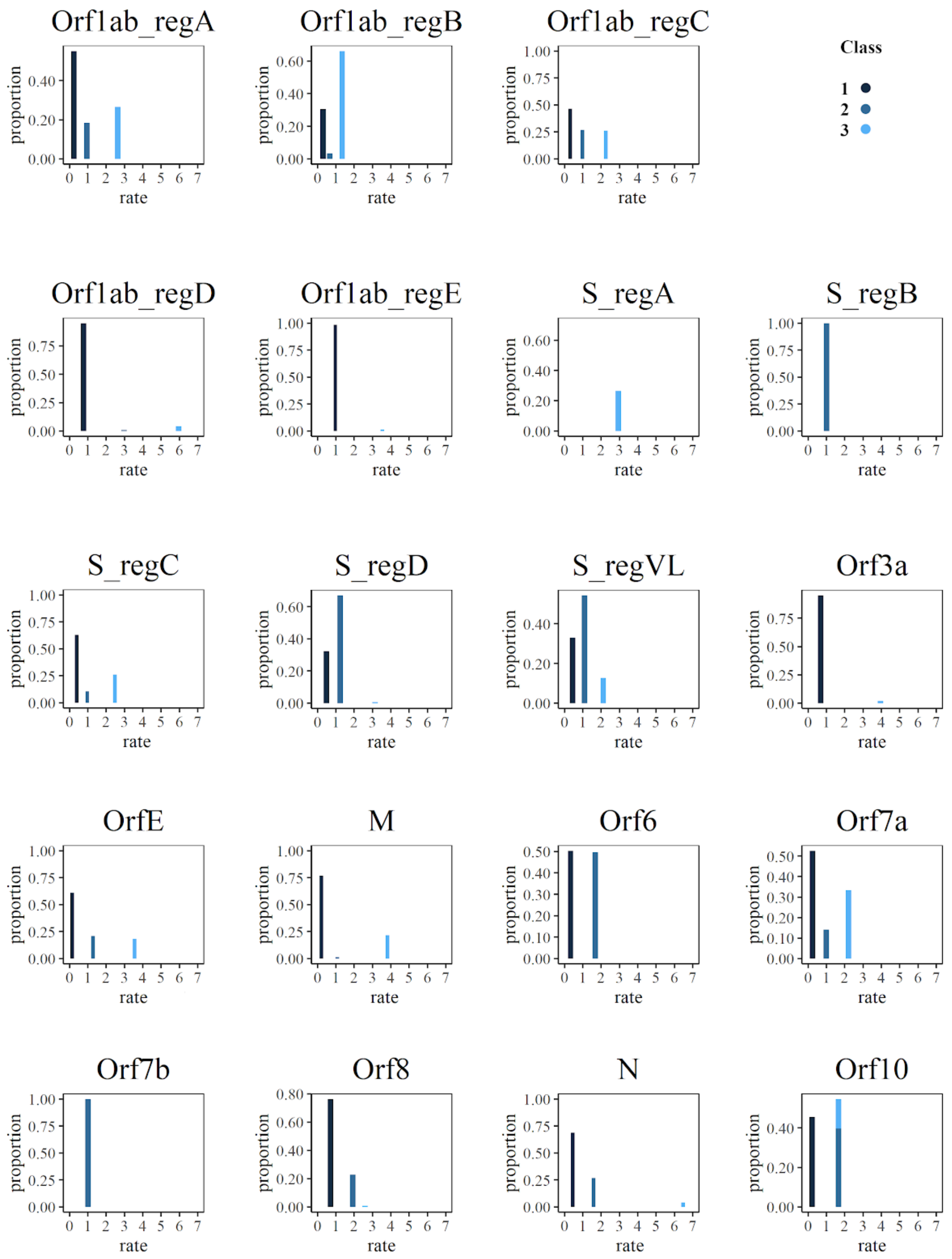


***Supplementary figure 6.*** *Synonymous rate of each of the three SRV classes estimated by BUSTED[S] + HMM and the proportion attributed to each class for all non-recombinant open reading frames.*

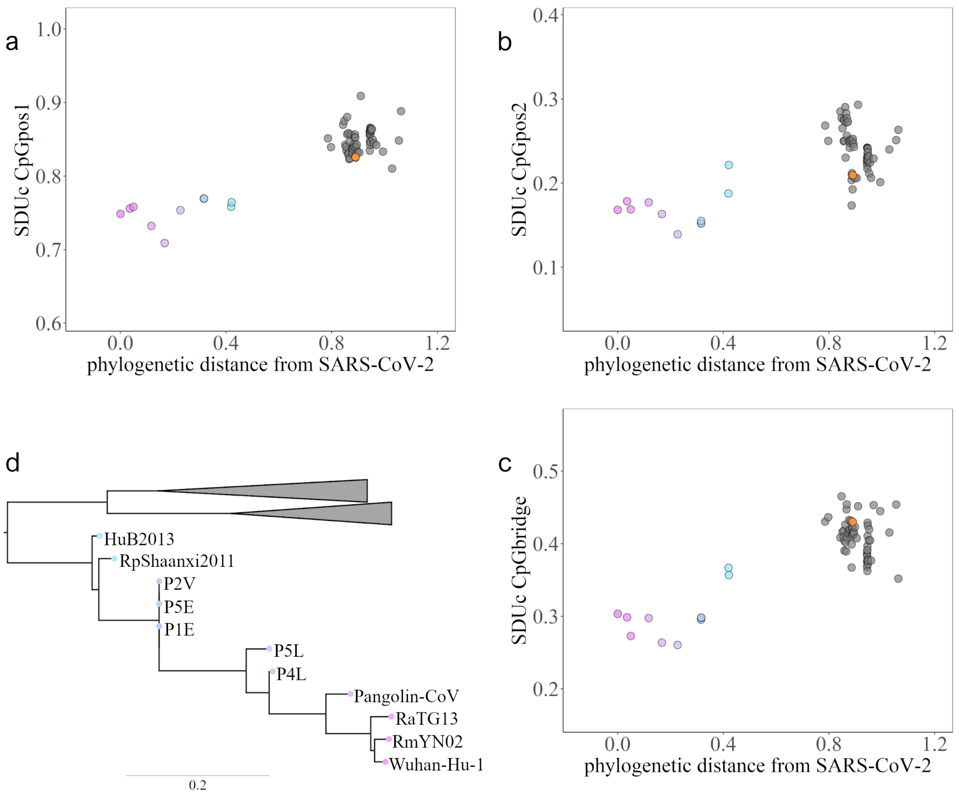


***Supplementary figure 7.*** *Corrected synonymous dinucleotide usage (SDUc) values for the Orf1ab of each Sarbecovirus for all dinucleotide frame positions (pos1 (a): 1st and 2nd codon positions; pos2 (b): 2nd and 3rd codon positions; bridge (c): 3rd codon position and 1st position of the next codon) plotted against patristic distance from SARS-CoV-2 (reference genome Wuhan-Hu-1). The tip colours of the phylogeny (d) correspond to the SDUc data points. SARS-CoV-1 is labelled in orange in a,b,c for comparison. The non-nCoV part of the phylogeny has been excised for clarity.*

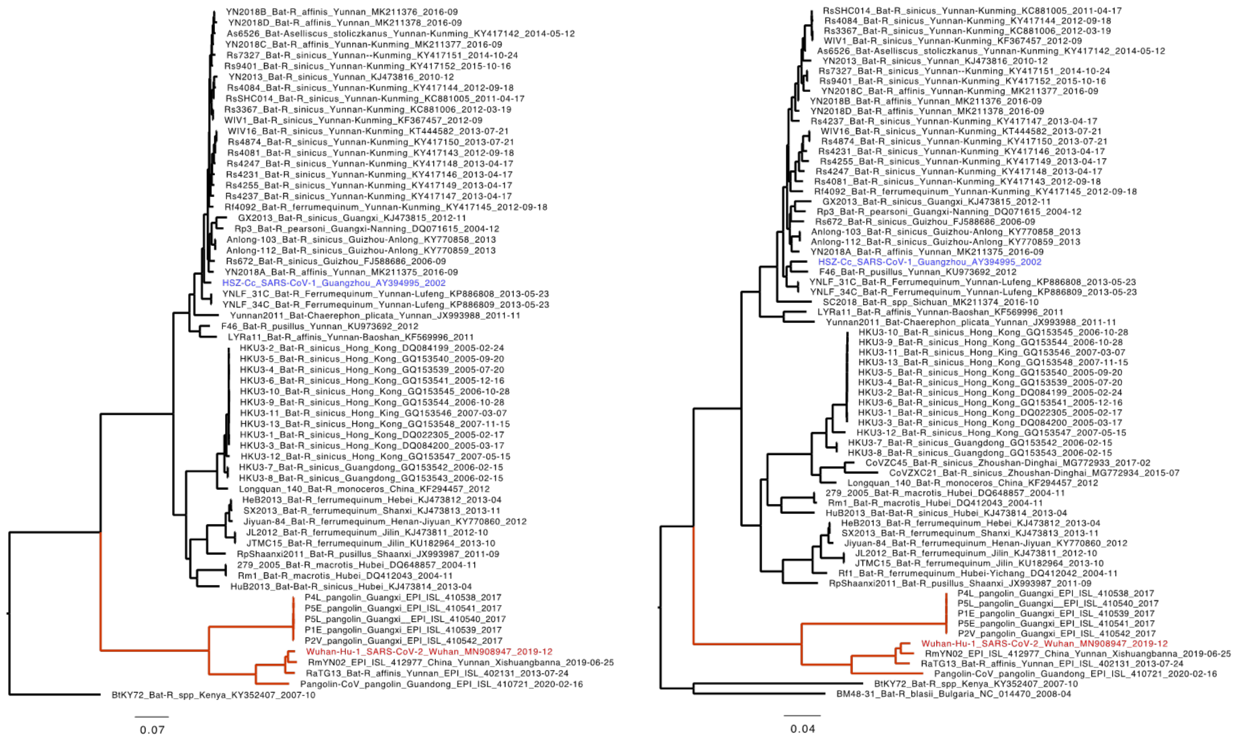

**Supplementary Figure 8.** Maximum likelihood trees for NRR1 (left) and NRR2 (right). The trees were inferred using IQTREE using a GTR substitution model with gamma-distributed rate variation among sites. The nCoV lineage is indicated in red. SARS and SARS-CoV-2 are shaded in blue and red respectively.

**Supplementary Table 5.** Mean divergence time estimates with 95% highest posterior density intervals for key nodes in the nCoV lineage.

| Divergence time | NRR1 | NRR2 |
|---|---|---|
| **nCoV** | 1189 (254,1538) | 1467 (1264,1641) |
| **Guangdong pangolin / SC2** | 1664 (1280,1815) | 1732 (1627,1825) |
| **RaTG13 /SC2** | 1924 (1810,1958) | 1955 (1932,1976) |
| **RmYN02 /SC2** | 1949 (1916,1975) | 1976 (1959,1991) |

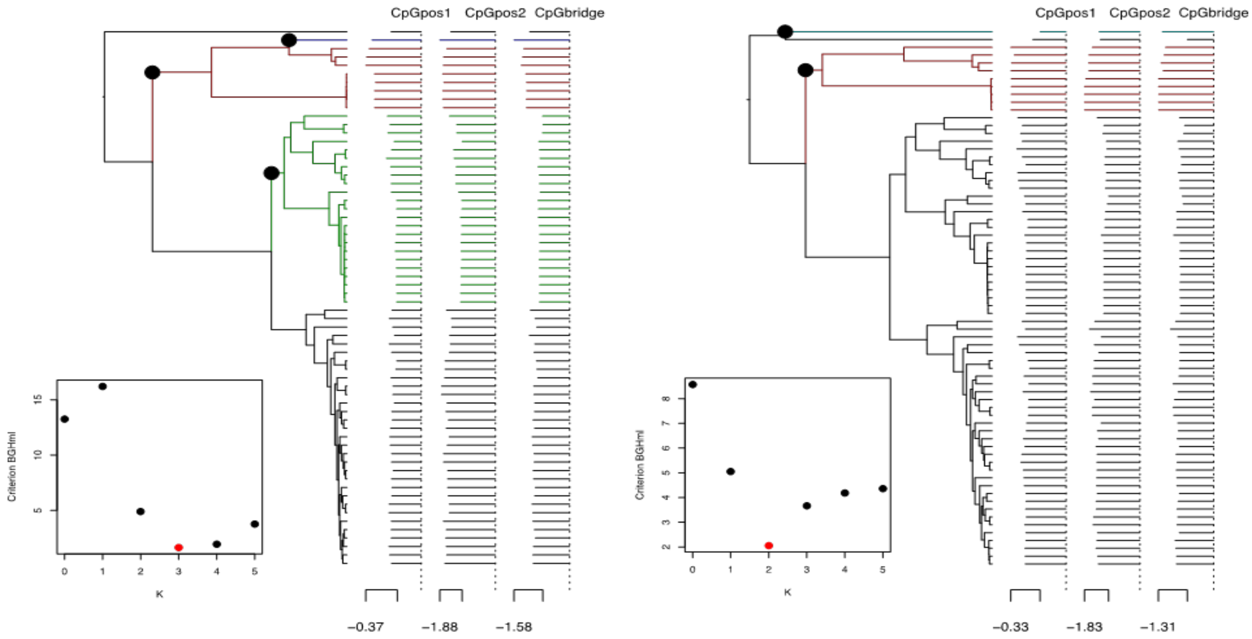


**Supplementary Figure 9.** Estimates of shifts in CpG in the Sarbecovirus phylogeny based on NRR1 (left) and NRR2 (right). The log-transformed CpG content values are shown at the tips of the trees. The identified shifts are indicated with black circles on their respective branches and with different colors for the lineages and CpG measures involved. The inset shows the results for penalized least-squares model selection criterion (BGHml)[14].

### Supplementary Text 4. Origin of the SARS-CoV-2 Spike furin cleavage site insertion

Contrary to all of its closely related *Sarbecoviruses*, SARS-CoV-2 possesses a furin cleavage site insertion on its Spike ORF of SARS-CoV-2[15] required for infection of certain cell types[20]. Although whether this has a significant functional importance for the virus in the presence of other proteases is controversial[16], it is still of scientific interest to determine how this insertion originated in this virus's genome.

We have estimated that ancestors of RmYN02 and SARS-CoV-2 diverged from one another decades ago (Main text). However, the RmYN02 genome has a long recombinant region encompassing the first half of the Spike ORF[17], acquired from a currently unsampled viral lineage. Part of this region is also where the sequence corresponding to the SARS-CoV-2 furin site is. Considering the period of time since RmYN02 and SARS-CoV-2 shared a common ancestor, and the clear recombination evidence between the first and the unknown virus (we will be referring to here as clade X virus), it is reasonable to assume that all three viruses co-circulated in the same bat population at some point in time, occasionally co-infecting the same individuals.

When aligning the furin site region between RmYN02 and SARS-CoV-2, alignment algorithms show the furin site sequence as the inserted region (Supplementary Figure 10). Interestingly, we noticed that there is higher nucleotide sequence identity (inferring homology) between the RmYN02 sequence and the inserted SARS-CoV-2 furin site region, despite the larger number of gaps under this alignment (compare Supplementary Figure 10A to 10B). Given that the clade X viruses recombined with SARS-CoV-2's sister lineage, RmYN02, we propose that the furin site insertion was acquired from a clade X virus when co-infecting the same bat individual. The exact molecular mechanism through which this insertion took place cannot currently be inferred, but a likely explanation would be a copy choice error of the viral polymerase or a short template switch during negative RNA synthesis[18,19]. The two single nucleotide gaps on the 3' of the furin site in each viral sequence would then be explained by slippage when the polymerase switched to the heterologous sequence. Six in-frame nucleotides encoding for two arginines (CGG CGG) are also missing from the RmYN02 genome (Supplementary Figure 10B). We suggest that these have been deleted in RmYN02 since the recombination event with the clade X virus.

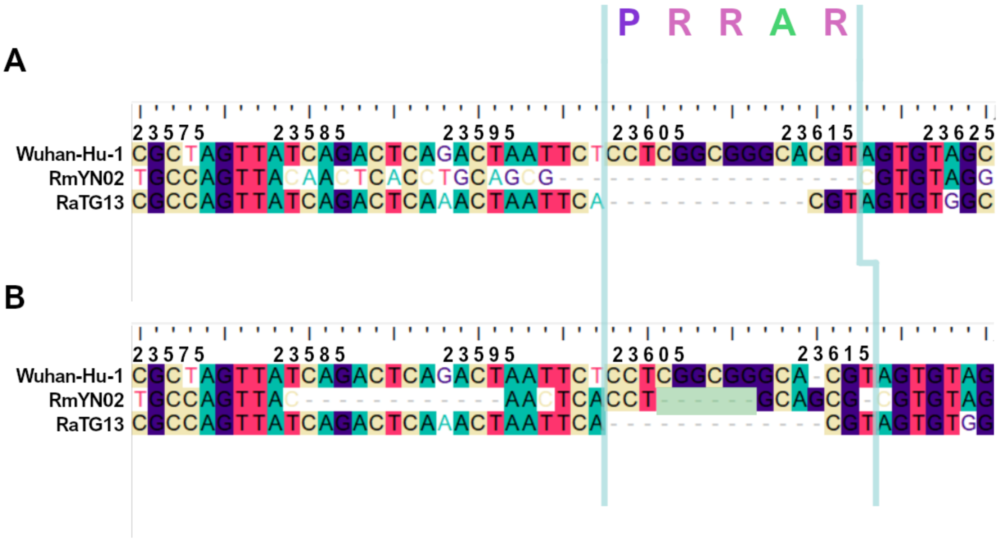


***Supplementary Figure 10.*** *Alternative alignments of the SARS-CoV-2 reference sequence (Wuhan-Hu-1) with RmYN02 at the furin site region. The low sequence identity from positions 23,585 to 23,599 between Wuhan-Hu-1 and RmYN02 (panel A), relative to the alternative alignment (panel B), shows there's a homologous region to the furin cleavage site present in RmYN02 (positions 23603 to 23615). This indicates the inserted region in SARS-CoV-2 progenitor was acquired by recombination in a host co-infected with divergent sarbecoviruses. RaTG13 is also included for clarity. Nucleotide coordinates are mapped to the Wuhan-Hu-1 whole genome sequence. The putative deletion of 6 nucleotides in RmYN02 are highlighted in green.*

### *External documents*

**Supplementary Table 1.** List of GISAID accessions used for FUBAR analysis.

**Supplementary Table 3.** List of sites found to be under selection by MEME in the nCoV clade.

**Supplementary Table 4.** List of ω distributions inferred for non-recombinant segments under BUSTED[S]-HMM.

**Supplementary Table 6.** List of *Sarbecoviruses* used in the selection analysis.

**Supplementary Table 7.** GISAID sequences acknowledgement table.
